# Structural polymorphism of the nucleic acids in pentanucleotide repeats associated with CANVAS

**DOI:** 10.1101/2023.08.17.553768

**Authors:** Kenta Kudo, Karin Hori, Sefan Asamitsu, Kohei Maeda, Yukari Aida, Mei Hokimoto, Kazuya Matsuo, Yasushi Yabuki, Norifumi Shioda

**Affiliations:** Department of Genomic Neurology, Institute of Molecular Embryology and Genetics (IMEG), Kumamoto University; Kumamoto 860-0811, Japan; Graduate School of Pharmaceutical Sciences, Kumamoto University; Kumamoto 862-0973, Japan; Laboratory for Functional Non-coding Genomics, RIKEN Center for Integrative Medical Sciences (IMS); Yokohama, 230-0045, Japan

**Keywords:** nucleic acid structure, biophysics, G-quadruplex, genetic disease, neurodegenerative disease

## Abstract

Short tandem repeats are highly unstable, depending on repeat length, and the expansion of the repeat length in the human genome is responsible for repeat expansion disorders. Pentanucleotide AAGGG and ACAGG repeat expansions in intron 2 of the gene encoding replication factor C subunit 1 (*RFC1*) cause cerebellar ataxia, neuropathy, vestibular areflexia syndrome (CANVAS) and other phenotypes of late-onset cerebellar ataxia. Herein, we reveal the structural polymorphism of the *RFC1* repeat sequences associated with CANVAS *in vitro*. Single-stranded AAGGG repeat DNA formed a hybrid-type G-quadruplex, whereas its RNA formed a parallel-type G-quadruplex with three layers. The RNA of the ACAGG repeat sequence formed double helical hairpin structures comprising C-G and G-C base pairs with A:A and GA:AG mismatched repeats. Furthermore, both pathogenic repeat RNAs formed more rigid structures than those of the non-pathogenic sequences. These findings provide novel insights into the structural polymorphism of the *RFC1* repeat sequences, which may be closely related to the disease mechanism of CANVAS.

## Introduction

Short tandem repeats (STRs) are polymorphic repeat units of 1–6 base pairs that are abundant in eukaryotic genomes [1]. STRs are extremely unstable, depending on repeat length, and their expansion in the coding and non-coding regions across generations is the cause of almost 50 repeat expansion disorders, most of which primarily affect the nervous system [2]. Thirteen STRs are known to cause repeat expansion disorders, and the structures of non-canonical DNA and its transcribed RNA, e.g., slipped hairpins, G-quadruplexes (G4s), and triplexes, that form within repeat regions may induce cytotoxicity via abnormal genomic stability, transcription, epigenetics, mRNA translation, and RNA transport and decay [3–5].

Cerebellar ataxia, neuropathy, vestibular areflexia syndrome (CANVAS) is an autosomal recessive, slowly progressive, late-onset neurological disorder clinically characterized by the triad of cerebellar ataxia, bilateral vestibular hypofunction, and a peripheral sensory deficit [6,7]. Recently, a biallelic AAGGG repeat expansion (AAGGG)_exp_ ranging from 400 to 2000 repeats in the gene encoding replication factor C subunit 1 (*RFC1*) was reported to cause CANVAS and other phenotypes of late-onset cerebellar ataxia [8,9]. In addition to the pathogenic (AAGGG)_exp_ allele, two other pathogenic alleles were identified in the same *RFC1* locus: (ACAGG)_exp_ in the Asia-Pacific and Japanese cohorts [10,11] and (AAAAG)_10–25_(AAGGG)_exp_(AAAGG)_4–6_ in the Māori-specific cohort [12]. Remarkably, three different repeat conformations are observed in non-pathogenic populations: the reference (AAAAG)_11_, (AAAAG)_exp_ ranging from 15 to 200 repeats, and (AAAGG)_exp_ ranging from 40 to 1000 repeats (**Fig. 1A**) [8]. The cumulative allele frequency of (AAAAG)_exp_ and (AAAGG)_exp_ of approximately 20 % in the general population is relatively high [8]. A positive correlation between repeat length and G/C content is observed, whereas no correlation between repeat length and age of onset exists [10,13]. Thus, the onset of CANVAS may be due to the secondary structures of nucleic acids formed by the G/C-rich composition, independent of the repeat length. However, there are no studies on the secondary structures of nucleic acids associated with CANVAS.

**Figure 1.**
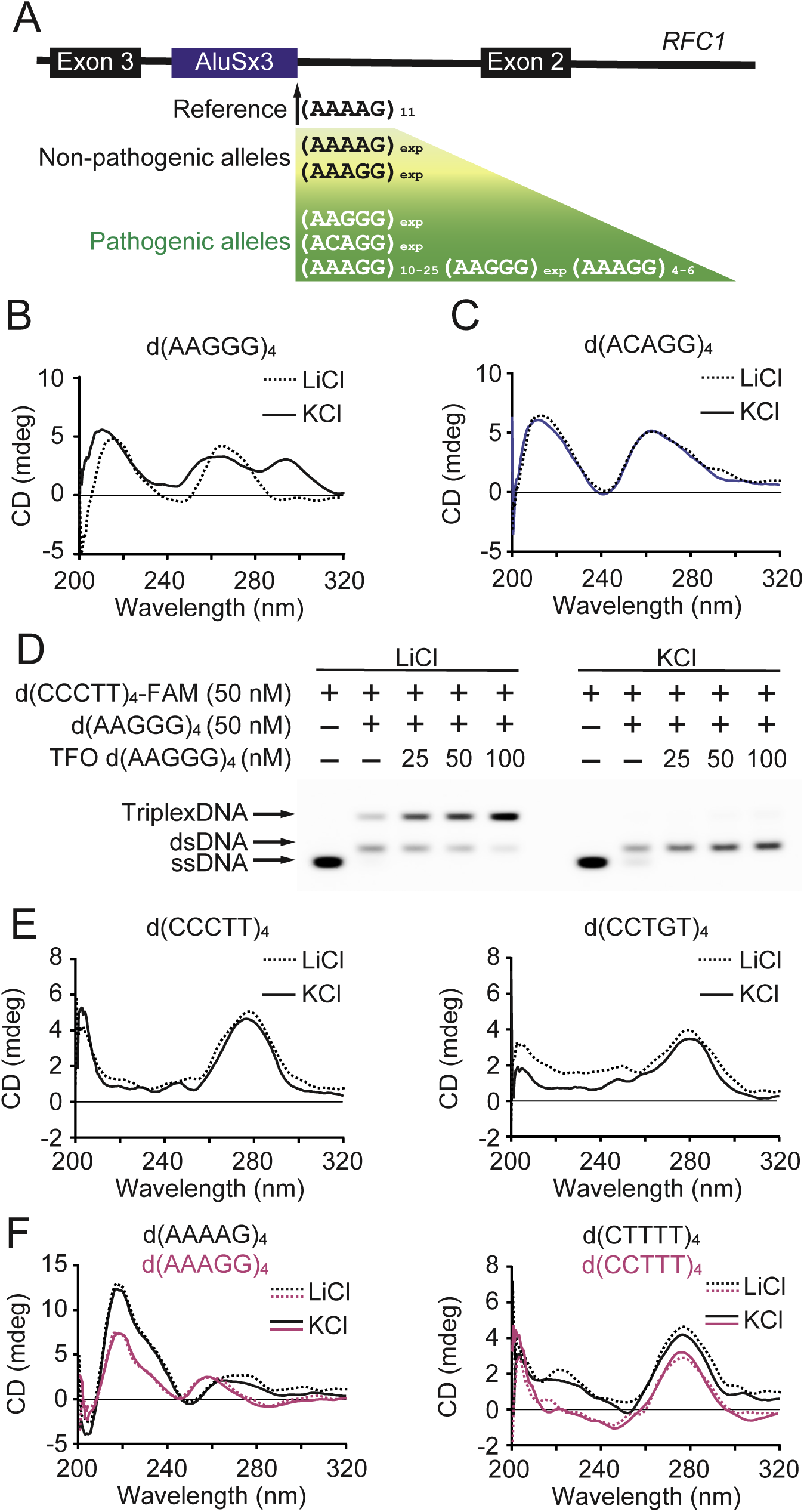
CD spectra of the DNAs in *RFC1* pentanucleotide repeats. (A) Schematic diagram showing the polymorphism at the *RFC1* repeat locus, consisting of non-pathogenic and pathogenic alleles. *RFC1*: replication factor C subunit 1. See also Davies et al., (2022). (B, C) Representative CD spectra of the pathogenic sequences d(AAGGG)_4_ and d(ACAGG)_4_ in the presence of 100 mM KCl (solid line) or LiCl (dotted line) at 25 °C and pH 7.5. (D) Gel mobility shift assays of a purine triplex at physiological pH 7.5 in the presence of Mg^2+^ and 100 mM LiCl (left) or KCl (right). (E) Representative CD spectra of the pathogenic complementary sequences d(CCCTT)_4_ and d(CCTGT)_4_ in the presence of 100 mM KCl (solid line) or LiCl (dotted line) at 25 °C and pH 7.5. (F) Representative CD spectra of non-pathogenic d(AAAAG)_4_ and d(CTTTT)_4_ (black lines) and d(AAAGG)_4_ and d(CCTTT)_4_ (magenta lines) in the presence of 100 mM KCl (solid line) or LiCl (dotted line) at 25 °C and pH 7.5. All experiments were repeated at least twice, and the results were similar.

Herein, we report the secondary structures of nucleic acids related to CANVAS using *in vitro* CD and UV-Vis spectroscopy and thioflavin T (ThT) fluorescence and footprinting assays. Additionally, based on the results, the higher-order structures of nucleic acids in pathogenic repeat sequences are simulated via molecular modeling. Single-stranded AAGGG repeat DNA and its RNA respectively form hybrid- and parallel-type G4s, and furthermore, ACAGG repeat RNA forms a unique slipped-hairpin structure. Conversely, none of the non-pathogenic repeat sequences form characteristic nucleic acid secondary structures. The disease-specific formation of non-canonical nucleic acid structures may be the underlying molecular cause of the pathology of CANVAS.

## Results

### CD of the DNAs in *RFC1* pentanucleotide repeats

We first investigated the topological unimolecular DNA structures of 20-mer single-stranded oligonucleotides of the non-pathological sequences d(AAAAG)_4_ and d(AAAGG)_4_, pathological sequences d(AAGGG)_4_ and d(ACAGG)_4_, and their complementary sequences d(CTTTT)_4_, d(CCTTT)_4_, d(CCCTT)_4_, and d(CCTGT)_4_ using CD. It was performed at physiological pH, i.e., pH 7.5.

Single-stranded guanine-rich sequences may form G4s, which are four-stranded nucleic acid structures comprising stacked G-quartets, i.e., planar arrays of four guanines stabilized by Hoogsteen base pairing involving the N at position 7 of guanine [14]. In the presence of 100 mM K^+^, which stabilizes G4s, d(AAGGG)_4_ forms a well-defined hybrid-type G4 structure [15], resulting in a CD spectrum containing positive peaks at approximately 295 and 260 nm. In contrast, in the presence of G4-non-stabilizing 100 mM Li^+^, the entire spectrum changes, and the typical features are not observed (**Fig. 1B**). To further confirm the G4 folding of d(AAGGG)_4_, we assessed the cation-dependent thermal denaturing by measuring the melting temperature (*T*m). d(AAGGG)_4_ exhibits a much higher thermal stability under K^+^ conditions than that under Li^+^ conditions (Δ*T*m = 10.4 °C). No characteristic spectrum of the other pathological sequence, d(ACAGG)_4_, is observed in the presence of K^+^ or Li^+^ (**Fig. 1C**).

Homopurine/-pyrimidine mirror repeats, such as AAGGG/CCCTT repeat sequences, may adopt intramolecular triplex structures [16]. Triplexes are classified into two types according to the base composition and binding direction of the triplex-forming oligonucleotide (TFO) against the duplex DNA: pyrimidine and purine triplexes. Pyrimidine triplexes comprise TA-T and protonated CG-C^+^ triplets, which require a weakly acidic environment for formation, whereas purine triplexes, which comprise CG-G and TA-A triplets, are stable at physiological pH in the presence of Mg^2+^ [17]. Herein, we investigated whether purine triplexes form at physiological pH 7.5 with Mg^2+^ using the gel shift assay. Using single-stranded d(AAGGG)_4_ as the TFO and fluorescein-labeled double-stranded d(AAGGG/CCCTT)_4_, a triplex is detected in the presence of Li^+^, depending on the TFO concentration (**Fig. 1D, left**). In the presence of K^+^, where the TFO single-stranded d(AAGGG)_4_ forms a G4, triplex formation is reduced compared to that under the Li^+^ conditions (**Fig. 1D, right**). A G4 may be more likely to form than a triplex under near-physiological conditions when K^+^ is present.

A cytosine-rich DNA sequence forms an i-motif, which is a structure comprising two intercalated parallel-stranded duplexes held together by hemi-protonated C:C^+^ base pairs [18]. However, the CD spectra of the complementary cytosine-rich single-stranded d(CCCTT)_4_ and d(CCTGT)_4_ display positive peaks at 275 nm (**Fig. 1E**). No CD profile features of the i-motif structure are observed, with positive and negative peaks at 288 and 264 nm, respectively [19].

In terms of the non-pathological sequences, the CD spectra of adenine-rich single-stranded d(AAAAG)_4_ and d(AAAGG)_4_ display strong positive peaks at 220 nm, and those of thymine-rich single-stranded d(CTTTT)_4_ and d(CCTTT)_4_ display positive peaks at 275 nm (**Fig. 1F**). Consistent with these results, previous studies report that the CD spectra of single-stranded poly(dA) and poly(dT) exhibit corresponding peaks at 220 and 275 nm, and correlations between these signal intensities and the total contents of adenine and thymine are observed, independent of the presence of secondary structures [19,20].

### G4 conformation of AAGGG repeat DNA

ThT binds preferentially to G4s and emits a strong fluorescence at a low background [21]. To confirm the possibility that the AAGGG repeat sequence forms a G4, we used DNA oligonucleotides to measure the intensity of ThT fluorescence in the presence of K^+^. Consistent with the result of CD (**Fig. 1B**), a drastic increase in ThT fluorescence is detected for d(AAGGG)_4_ compared to those of other sequences (**Fig. 2A**).

**Figure 2.**
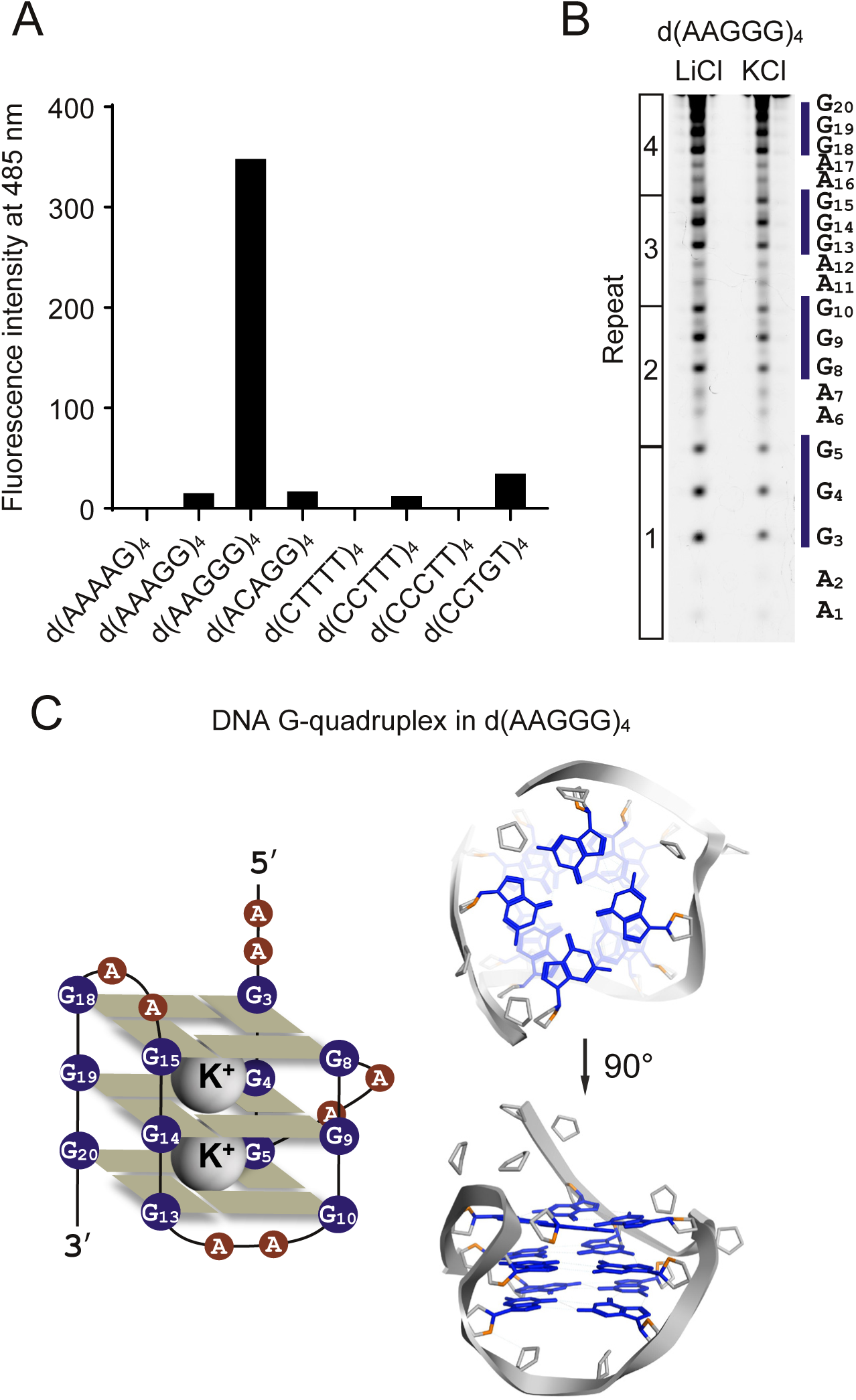
G4 conformation of AAGGG repeat DNA. (A) Bar graph of the fluorescence intensities of ThT at 485 nm in the presence of various DNA oligonucleotides. Samples were prepared at 2.5 µM in 10 mM Tris/HCl at pH 7.5 with 100 mM KCl, and ThT was added to yield a concentration of 0.2 µM. Experiments were repeated at least twice, and the results were similar. (B) DMS footprinting of d(AAGGG)_4_ oligonucleotides in the presence of 100 mM K^+^ or Li^+^. The sequence of d(AAGGG)_4_ is shown to the right of the gel. Experiments were repeated at least twice, and the results were similar. (C) Schematic and 3D diagrams of the G4 conformation of d(AAGGG)_4_, based on the results of the biophysical analyses.

To elucidate the G4 formation of d(AAGGG)_4_ in detail, we performed dimethyl sulfate (DMS) footprinting and identified the guanines involved in the G-quartets based on their resistances to methylation at position N^7^ due to Hoogsteen hydrogen bonding. In the presence of K^+^, all guanines (G3–G5, G8–G10, G13–G15, and G18–G20) are more sufficiently protected against DMS methylation and subsequent piperidine cleavage than in the presence of Li^+^ (**Fig. 2B**). The results of CD (**Fig. 1B**) and the ThT assay (**Fig. 2A**), in addition to the DMS footprinting pattern, suggest G4 formation by d(AAGGG)_4_, where 4 tracts of 3 guanines form 4 strands with 3 two-base loops comprising AA (**Fig. 2C**).

### CD of the RNAs in *RFC1* pentanucleotide repeats

Similar to CD of the DNAs, the topological unimolecular RNA structures were identified using CD under physiological conditions at pH 7.5 and oligonucleotides of the non-pathological and pathological sequences and their complementary sequences.

In the presence of K^+^, r(AAGGG)_4_ forms a typical parallel-stranded G4, exhibiting positive and small negative peaks at 262 and 240 nm, respectively, and a positive band at 210 nm [15], which is not observed under Li^+^ conditions (**Fig. 3A, left**). To investigate the higher-order RNA structures with more pathological repeat lengths, we performed CD using r(AAGGG)_11_, and the spectrum displays a peak more characteristic of a parallel-type G4 compared to that in the spectrum of r(AAGGG)_4_ (**Fig. 3A, right**). In the thermal denaturation assay at the same concentration, the stabilizing effects of K^+^ on r(AAGGG)_11_ (Δ*T*m = 41.0 °C) and r(AAGGG)_4_ (Δ*T*m = 44.0 °C) compared to those of Li^+^ are observed. In the CD spectrum of ACAGG repeat RNA, no characteristic peaks representing r(ACAGG)_4_ are observed. Conversely, the positive and negative CD absorption bands of r(AAGGG)_11_ are detected as a maximum at ∼260 nm, a minimum at ∼210 nm, and a small negative peak between 290 and 300 nm under Li^+^ and K^+^ conditions (**Fig. 3B**). These peaks are characteristic of the A-form duplex RNA conformation [22].

**Figure 3.**
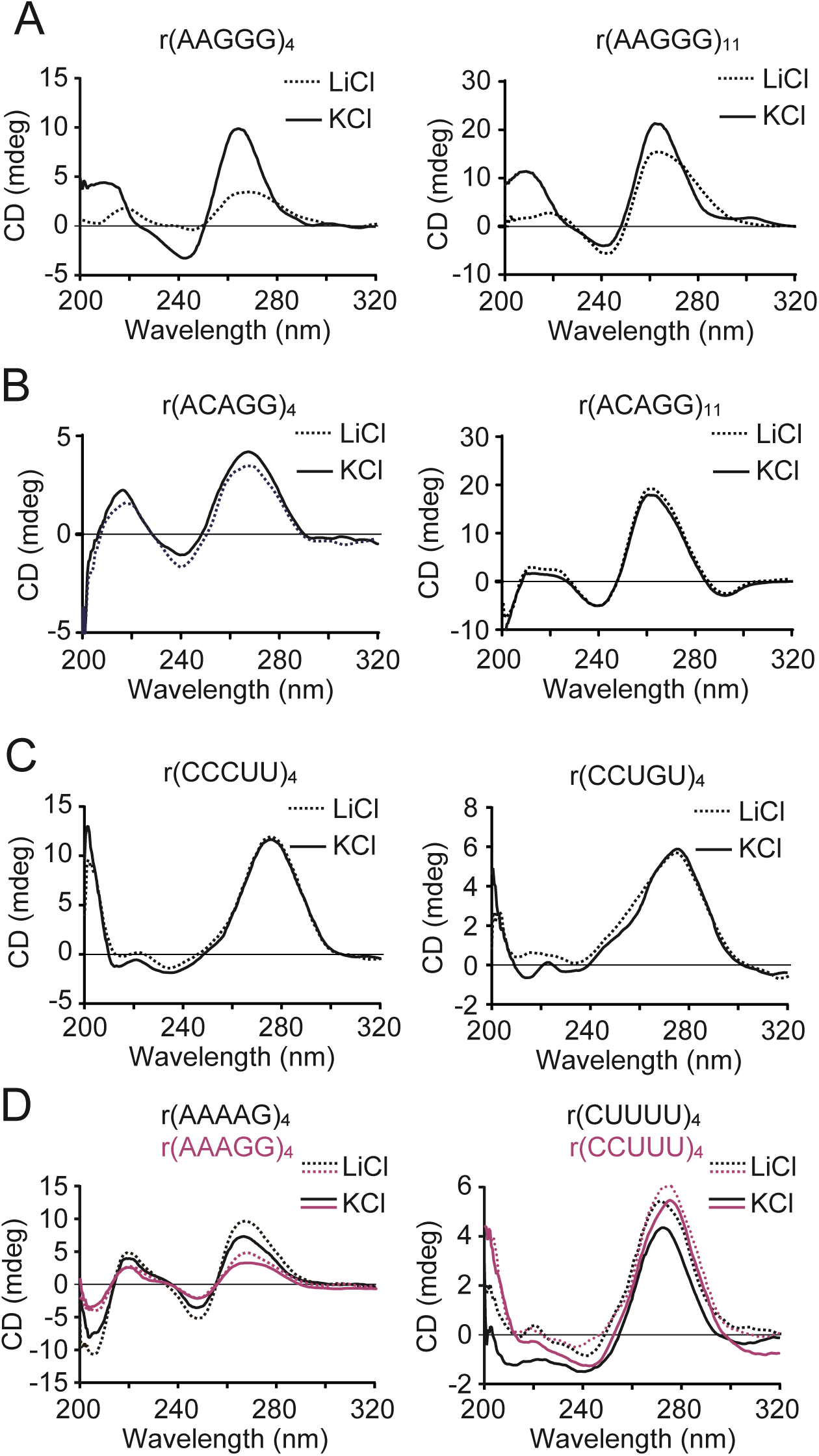
CD spectra of the RNAs in *RFC1*-derived pentanucleotide repeats. (A, B) Representative CD spectra of pathogenic r(AAGGG)_4_, r(AAGGG)_11_, r(ACAGG)_4_, and r(ACAGG)_11_ in the presence of 100 mM KCl (solid lines) or LiCl (dotted lines) at 25 °C and pH 7.5. (C) Representative CD spectra of the pathogenic complementary sequences r(CCCUU)_4_ and r(CCUGU)_4_ in the presence of 100 mM KCl (solid lines) or LiCl (dotted lines) at 25 °C and pH 7.5. (D) Representative CD spectra of non-pathogenic r(AAAAG)_4_ and r(CUUUU)_4_ (black lines) and r(AAAGG)_4_ and r(CCUUU)_4_ (magenta lines) in the presence of 100 mM KCl (solid line) or LiCl (dotted line) at 25 °C and pH 7.5. All experiments were repeated at least twice, and the results were similar.

None of the CD spectra of the complementary strands of the pathological sequences, r(CCCUU)_4_ and r(CCUGU)_4_, display characteristic peaks indicative of RNA secondary structures (**Fig. 3C**). In terms of the non-pathological sequences, the CD spectra of the adenine-rich single strands r(AAAAG)_4_ and r(AAAGG)_4_ display strong positive and small negative peaks at 265 and 250 nm, respectively (**Fig. 3D, left**), which are consistent with the tendency of poly(A), based on its CD spectrum, to form unique single-stranded helical structures [23]. The CD spectra of uridine-rich single-stranded d(CUUUU)_4_ and d(CCUUU)_4_ display small positive peaks at 275 nm (**Fig. 3D, right**), indicating the absence of complex structures, similar to poly(U) [23].

### Analyses of the RNA structures of the pathological repeat sequences related to CANVAS

The intensity of ThT fluorescence was then measured to investigate the possibility of G4 formation using 20-mer RNAs with non-pathological and pathological sequences and their complementary sequences. A drastic increase in ThT fluorescence is observed for r(AAGGG)_4_ compared to those of the other repeat sequences (**Fig. 4A**).

**Figure 4.**
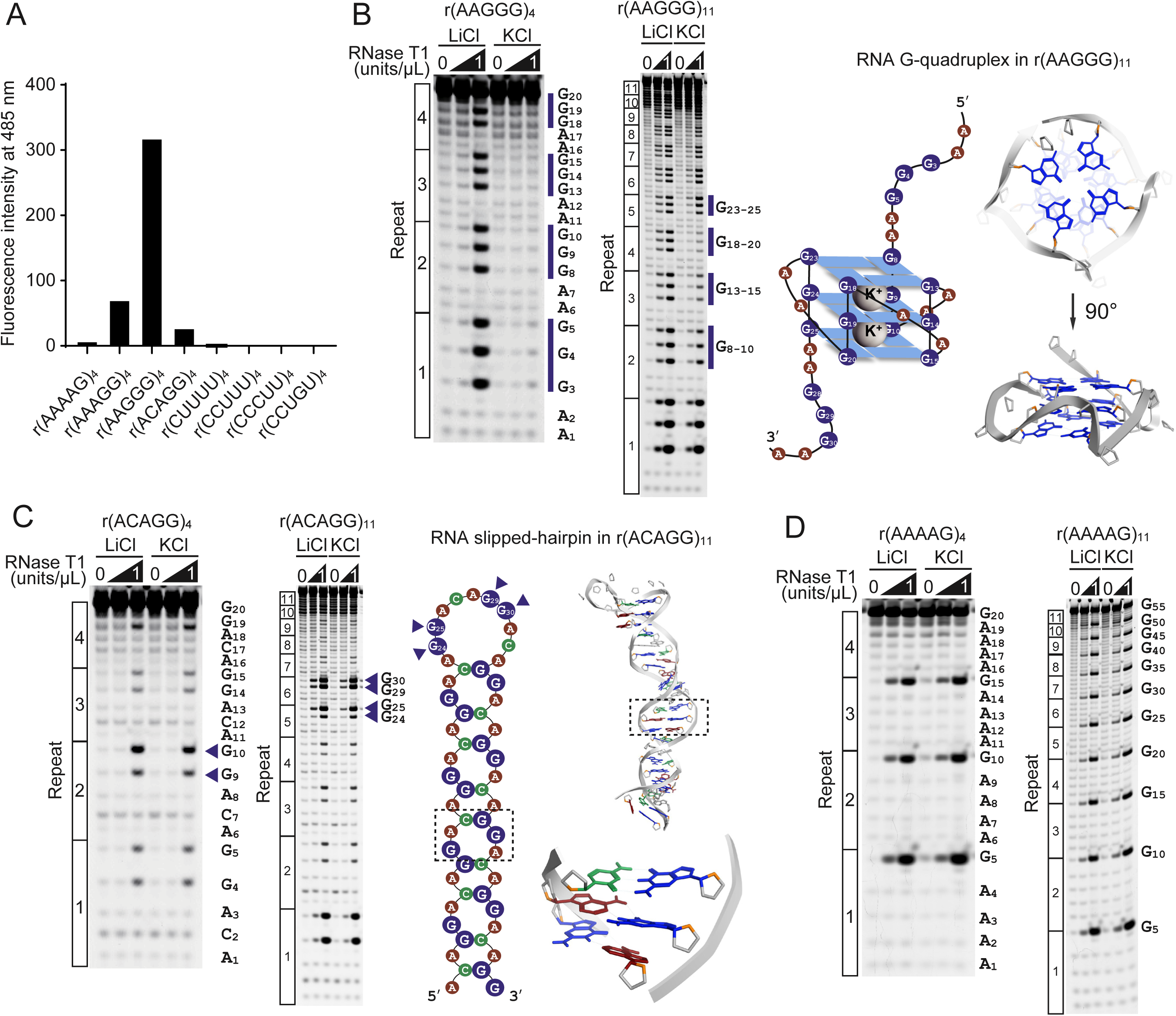
Secondary structures of the pathogenic repeat RNAs related to CANVAS. (A) Bar graph of the fluorescence intensities of ThT at 485 nm in the presence of various RNA oligonucleotides. Samples were prepared at 2.5 µM in 10 mM Tris/HCl at pH 7.5 with 100 mM KCl, and ThT was added to yield a concentration of 0.2 µM. (B) Patterns of RNase T1 cleavage of r(AAGGG)_4_ (left) and r(AAGGG)_11_ (middle). The vertical blue lines show the differences in the cleavage patterns obtained in the presence of LiCl and KCl. Schematic and 3D diagrams of the G4 conformation of r(AAGGG)_4_, based on the results of the biophysical analyses (right). (C) Patterns of RNase T1 cleavage of r(ACAGG)_4_ (left) and r(ACAGG)_11_ (middle). The blue triangles indicate strongly truncated guanosines that may be involved in loop formation. Schematic and 3D diagrams of the hairpin conformation of r(ACAGG)_4_, based on the results of the biophysical analyses. The G-C and G:A base pairs are putatively formed (right). (D) Patterns of RNase T1 cleavage of r(AAAAG)_4_ (left) and r(AAAAG)_11_ (right). All experiments were repeated at least twice, and the results were similar.

To identify more detailed RNA structures, we conducted RNase T1 footprinting of pathological AAGGG and ACAGG and non-pathological AAAAG repeats. As RNase T1 displays specificity for guanine residues in single-stranded RNA, guanosines in loops and unstructured regions are more susceptible to cleavage than those directly involved in G4 and hairpin formation [24]. In r(AAGGG)_4_, all twelve guanosines (G3–G5, G8–G10, G13–G15, G18–G20) are protected against RNase T1 cleavage in the presence of K^+^ (**Fig. 4B, left**). Based on the RNase footprinting pattern of r(AAGGG)_11_ (**Fig. 4B, middle**) and the results of CD (**Fig. 3A**), and the ThT assay (**Fig. 4A**), the pathological AAGGG repeat sequence displays a strong tendency to fold into an intramolecular parallel-form G4, which comprises 3 G-quartets with 3 two-base loops comprising AA (**Fig. 4B, right**). Conversely, the guanosines in r(ACAGG)_4_ are cleaved, regardless of the monovalent cation added, and the intensities of guanosines G9 and G10 are slightly increased in the cleavage pattern (**Fig. 4C, left**). In the RNase footprinting pattern of r(ACAGG)_11_, four guanosines (G24, G25, G29, G30) predicted to occur in the single-stranded loop region are cleaved by RNase T1 (**Fig. 4C, middle**). RNA secondary structure prediction using deep learning based on MXfold2 (Keio University, Yokohama, Japan) [25] reveals that the stem structure formed by repeats of ACAGG on the 5′ and 3′ sides comprises C-G and G-C base pairs and A:A and GA:AG mismatch repeats (**Fig. 4C, right**). This is consistent with the results obtained, including those of CD (**Fig. 3B**). The energy-minimized structure of the r(ACAGG)_11_ hairpin conformation suggests that the hydrogen bonds of the G-C and G:A base pairs contribute to the stability of the structure, whereas the A:A mismatch does not form hydrogen bonds (**Fig. 4C, right**). In the non-pathological AAAAG repeat RNA, the single guanosine in the repeat sequence is cleaved by RNase T1 to the same extent in 4 and 11 repeats, indicating that r(AAAAG)_exp_ may not form a characteristic secondary structure (**Fig. 4D**).

To determine the differences in the structural stabilities of these RNAs, we performed UV-monitored melting studies. The obtained results confirm that r(AAGGG)_11_ and r(ACAGG)_11_ form stable structures, with respective changes in folding free energy of ΔG37°C = −24.44 ± 7.42 and −1.72 ± 0.11 kcal/mol, whereas that of r(AAAAG)_11_ is not detected (**Table 1**).

**Table 1.**
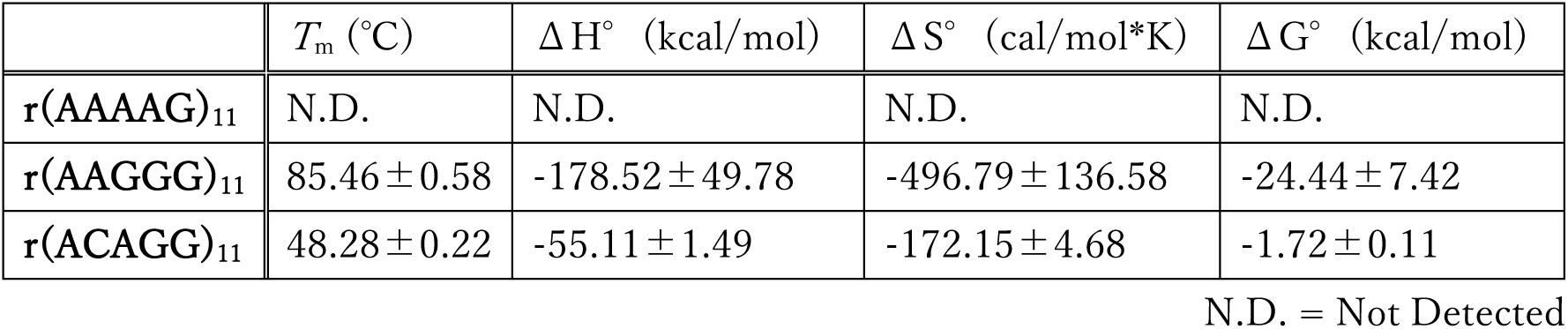
Experimentally obtained thermodynamic parameters of the RNA sequences. Experiments were repeated thrice.

**Figure 4**
| | $T_m$ (°C) | $\Delta H^\circ$ (kcal/mol) | $\Delta S^\circ$ (cal/mol*K) | $\Delta G^\circ$ (kcal/mol) |
| --- | --- | --- | --- | --- |
| **r(AAAAG)<sub>11</sub>** | N.D. | N.D. | N.D. | N.D. |
| **r(AAGGG)<sub>11</sub>** | $85.46 \pm 0.58$ | $-178.52 \pm 49.78$ | $-496.79 \pm 136.58$ | $-24.44 \pm 7.42$ |
| **r(ACAGG)<sub>11</sub>** | $48.28 \pm 0.22$ | $-55.11 \pm 1.49$ | $-172.15 \pm 4.68$ | $-1.72 \pm 0.11$ |

N.D. = Not Detected

## Discussion

We demonstrated the structural polymorphism of the *RFC1* repeat sequences associated with CANVAS. The single-stranded AAGGG repeat DNA and its RNA respectively formed hybrid- and parallel-type G4s, and the RNA of the ACAGG repeat formed a unique slipped-hairpin structure. Furthermore, both pathogenic repeat RNAs formed rigid structures compared to those of the non-pathogenic repeat sequences. These novel findings regarding structural polymorphism in *RFC1* repeat extensions may be relevant to the disease mechanism of CANVAS.

As G4s are formed by the stacking of at least two G-quartets, one possibility was that the ACAGG repeat sequences in DNA and RNA could also form G4s based on the consensus motif (G_2–3_ + N_1–7_G_2–3_ + N_1–7_G_2–3_ + N_1–7_G_2–3_) [4,26]. However, ACAGG repeats in RNA could not form G4s, instead forming unique slipped hairpin conformations, with the C-G and G-C base pairs combined with the A:A and GA:AG mismatched pairs. This is likely because G4 stability depends on several factors, including the lengths of the loops, its base composition, and the number of G tracts [27]. The stability of a G4 in RNA decreases with increasing loop length, particularly with two stacked G-quartets with the length of each loop longer than three residues, which is extremely unstable [28]. Additionally, the presence of cytosine relative to adenine and uridine leads to instability in a G4 with two G-quartets, because G-C Watson-Crick base pairs [29] predominate at high cytosine contents in the loops [28]. Furthermore, in ACAGG repeat RNA, the double helical structure of the A form may manifest, with sheared G:A and A:G base pairing in the loops [30,31], similar to that of the analogous base pairs in hammerhead ribozymes [32]. This study suggests that the combination of these factors results in ACAGG repeats with high energies, forming a rigid secondary structure not observed with non-pathogenic repeats.

We showed that neither single-stranded d(AAGGG)_4_ with double-stranded d(AAGGG/TTCCC)_4_ nor complementary pyrimidine-rich single-stranded d(CCCTT)_4_ formed triplex and i-motif structures under physiological conditions in the presence of K^+^. However, due to technical difficulties in preparing long single-stranded DNA, the repeat length-dependent structure could not be fully investigated. The formation of nucleic acid structures also varies depending on conditions such as molecular crowding and pH, and the stabilities of DNA in a triplex and i-motif vary with oligonucleotide length and acidic pH [33,34]. Investigating the formation of these non-canonical DNA structures with various repeat sizes under different conditions is necessary.

The structures of non-canonical DNA and its transcribed RNA that form during repeat expansion disorders are central to the molecular mechanisms of toxicity via loss- or gain-of-function, RNA-binding protein (RBP) sequestration, and repeat-associated non-AUG (RAN) translation [5,35]. In C9orf72 amyotrophic lateral sclerosis and frontotemporal dementia (FTD) caused by GGGGCC repeat expansions (d(G_4_C_2_)_exp_) in the *C9orf72* gene [36,37], its transcribed r(G_4_C_2_)_exp_ forms G4s and disrupts neuronal function via various mechanisms by generating nuclear foci [38]. Additionally, sense r(G_4_C_2_)_exp_ and antisense r(C_4_G_2_)_exp_ transcribed RNAs are translated in all reading frames into dipeptide repeat proteins via RAN translation [39–41], resulting in these RAN proteins forming toxic aggregates [42–44]. In another pathological mechanism, G4-forming d(G_4_C_2_)_exp_ inhibits transcription, causing loss of function by reducing C9orf72 protein levels [42]. Fragile X-associated tremor/ataxia syndrome (FXTAS) is a neurodegenerative disease caused by d(CGG)_exp_ in the *FMR1* gene [45]. Transcribed r(CGG)_exp_ interacts with several RBPs that form RNA foci and induce cellular dysfunction [46]. Transcribed r(CGG)_exp_ also produces the toxic protein FMRpolyG via RAN translation [47], and G4-forming r(CGG)_exp_ induces the liquid-solid phase transition of FMRpolyG, causing neurological dysfunction [48]. Slipped-hairpin-forming d(CAG/CTG)_exp_ is a particularly common cause of repeat expansion disease [2]. Transcribed r(CAG)_exp_ within coding genes is translated into polyglutamine (polyQ) tracts, inducing polyQ toxicity [49]. Transcribed r(CAG)_exp_ and r(CUG)_exp_ adopt highly stable hairpin structures that form nuclear foci and induce RNA toxicity [3], and the transcripts may also induce cell death via RAN translation [39,50]. Additionally, r(CAG)_exp_-derived RAN translation products co-aggregate with r(CAG)_exp_ foci within the cytoplasm [51].

Currently, how the repeat expansions in the gene encoding *RFC1* contribute to the pathogenesis of CANVAS is unknown. To date, no reductions in *RFC1* mRNA and protein levels have been observed in peripheral tissues and postmortem brain samples from patients with biallelic (AAGGG)_exp_ in the *RFC1* gene. Additionally, no sense or antisense RNA foci were detected in the cerebellar vermes of patients. Conversely, *RFC1* pre-mRNA expression displayed increased intron 2 retention in the cerebella, frontal cortices, lymphoblasts, and muscles of patients compared to that of healthy controls [8]. GC-rich intron expansion, which is associated with intron retention, may inhibit splicing by altering RNA structure and/or splicing factor access to intron regulatory regions [52]. These results were obtained using a few samples, and thus, further studies are necessary to obtain accurate results regarding the transcription and protein translation of *RFC1*.

In conclusion, we identified the DNA and RNA secondary structures of pathological repeats associated with CANVAS *in vitro*. In the pathological sequences, AAGGG repeat DNA and RNA formed G4s, and ACAGG repeat RNA formed a unique slipped-hairpin structure. These disease-specific rigid structures may aid in elucidating the pathological mechanism of CANVAS.

### Experimental procedures Oligonucleotides

The following DNA and RNA oligonucleotides were synthesized by Hokkaido System Science (Sapporo, Japan): d(AAAAG)_4_, d(AAAGG)_4_, d(AAGGG)_4_, d(ACAGG)_4_, d(CTTTT)_4_, d(CCTTT)_4_, d(CCCTT)_4_, d(CCTGT)_4_, 5ʹ 6-carboxyfluorescein (FAM)-d(CCCTT)_4_, r(AAAAG)_4_, r(AAAGG)_4_, r(AAGGG)_4_, r(ACAGG)_4_, r(CTTTT)_4_, r(CCTTT)_4_, r(CCCTT)_4_, r(CCTGT)_4_, 5ʹ 6-FAM-r(AAAAG)_4_, 5ʹ 6-FAM-r(AAGGG)_4_, and 5ʹ 6-FAM-r(ACAGG)_4_; and Ajinomoto Bio-Pharma Services (Ibaraki, Japan): 5ʹ 6-FAM-r(AAAAG)_11_, 5ʹ 6-FAM-r(AAGGG)_11_, and 5ʹ 6-FAM-r(ACAGG)_11_.

### CD

Oligonucleotides (2.5 µM) were prepared in 10 mM Tris/HCl (pH 7.5) buffer containing 100 mM KCl or LiCl. The oligonucleotides were heated at 95 °C and gradually cooled to room temperature over 1.5 h prior to use. CD was conducted at 25 °C in the range 200– 400 nm using a J-1100 spectrometer (JASCO, Tokyo, Japan) and a quartz cuvette with a 3 mm path length. In the denaturing assays, the CD signal of each oligonucleotide at 265 (RNA) or 296 nm (DNA) was monitored. Temperature scans were performed continuously from 20 to 95 °C at 1 °C/min, and the *T*_m_ values were determined as the temperature at half of the maximum decrease in the signal.

### ThT assays

Oligonucleotides (2.5 µM) were prepared in 10 mM Tris/HCl (pH 7.5) containing 100 mM KCl, and they were heated at 95 °C and gradually cooled to room temperature over 1.5 h prior to use. ThT (0.2 µM) was added to each sample, which was then incubated for 2 h at room temperature using a THERMO-SHAKER (Bio-Medical Science, Tokyo, Japan). The fluorescence spectra were recorded using a quartz cuvette with a 3 mm path length and an FP-8300 spectrometer (JASCO) at an excitation wavelength of 440 nm in the wavelength range 420–600 nm. The fluorescence signals of each oligonucleotide at 485 nm were used for comparison.

### Gel shift assays

d(AAGGG)_4_ and FAM-labeled d(CCCTT)_4_ were prepared in 20 mM Tris/HCl (pH 7.5) containing 20 mM MgCl_2_, 100 (or 0) mM LiCl, 100 (or 0) mM KCl, 2.5 mM spermidine, and d(AAGGG)_4_ (TFO) in the same conditions. The oligonucleotides were heated at 95 °C and gradually cooled to room temperature over 1.5 h prior to use. FAM-labeled duplex DNA (50 nM) was incubated with increasing concentrations of d(AAGGG)_4_ (TFO) in the buffer for 12 h at 37 °C. Electrophoresis was performed at 4 °C with TBE-urea polyacrylamide (29:1 acrylamide:bisacrylamide) (w/v) gel containing 20 mM MgCl_2_ at 100 V for 3 h in 1 × TBE running buffer. DNAs labeled with FAM on the gel were visualized using an Amersham TYPHOON (Cytiva, Marlborough, MA, USA).

### DMS footprinting

FAM-labeled DNA (2.5 µM) was prepared in 10 mM Tris/HCl (pH 7.5) containing 100 mM KCl or LiCl. The DNA oligomers were heated at 95 °C and gradually cooled to room temperature over 1.5 h prior to use. Each sample was incubated with 10 % DMS in 50 % ethanol for 1 min on ice. The reaction was stopped with a stop solution (0.1 M mercaptoethanol, 0.5 M sodium acetate, and 50 μg/mL calf thymus DNA). Samples purified via isopropanol precipitation were incubated with 10 % piperidine at 95 °C for 5 min and then dried using a CC-105 centrifugal concentrator (TOMY Digital Biology, Tokyo, Japan). The resulting residues were washed with RNase-free H_2_O, re-dried, and dissolved in a loading solution (80 % formamide and 10 mM NaOH). Aliquots of the resultant solution were loaded onto a 15 % TBE-urea polyacrylamide (19:1 acrylamide:bisacrylamide) (w/v) gel and electrophoresed at 1500 V for 2 h in 1 × TBE running buffer. DNAs labeled with FAM on the gels were visualized using the Amersham TYPHOON (Cytiva).

### RNase T1 footprinting

FAM-labeled RNA oligomers (1.5 µM) were prepared in 10 mM Tris/HCl (pH 7.5) buffer containing 100 mM KCl or LiCl. The RNA oligomers were heated at 95 °C and gradually cooled to room temperature over 1.5 h prior to use. The RNA was treated with RNase T1 (Thermo Fisher Scientific, Waltham, MA, USA) at room temperature for 3 min, and then a stop solution (8 M urea, 0.4 M 2-mercaptoethanol, and 0.5 M EDTA (pH 8.0)) was added to terminate the reaction. Aliquots of the resultant solution were loaded onto a 15 % TBE-urea polyacrylamide (19:1 acrylamide:bisacrylamide) (w/v) gel and electrophoresed at 1500 V for 2 h in 1 × TBE running buffer. The RNAs labeled with FAM on the gels were visualized using the Amersham TYPHOON (Cytiva).

### UV-vis spectral assays

RNA oligonucleotides (2.5 µM) were prepared in 10 mM Tris/HCl (pH 7.5) buffer containing 100 mM KCl. The RNA oligonucleotides were heated at 95 °C and gradually cooled to room temperature over 1.5 h prior to use. The absorbance of each RNA oligonucleotide at 260 (hairpin) or 295 nm (G4) was recorded using a V-750 spectrometer (JASCO) in a series of cell cuvettes with path lengths of 10 mm. Temperature scans were performed continuously from 20 to 95 °C at 1 °C/min, and the *T*_m_ values were determined using a 2-point method in the Melting Analysis program of the Spectra Manager software (JASCO) [53].

### Thermodynamic analysis

The thermodynamic parameters of the hairpin and G4 structures were determined according to a previously reported method [53,54]. The folding (*A*_*F*_(*T*)) and unfolding (*A*_*U*_(*T*)) baselines were manually selected in Excel (Microsoft, Redmond, WA, USA), where T represents the temperature in degrees Celsius.

Using the baselines, the equations representing the folded fractions (*N*_T_) are as follows:

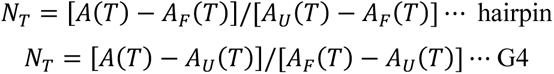

Under the two-transition-state assumption, the association constants (*K*_a_) are related to the folded fractions according to the following equations:

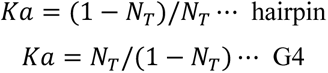

The Gibbs free enthalpy may be written as:

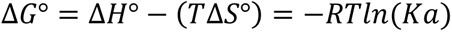

Where T, Δ*H*°, Δ*S*°, and *R* are the temperature in Kelvin, standard enthalpy of the reaction, standard entropy, and ideal gas constant, respectively.

The following van’t Hoff equation,

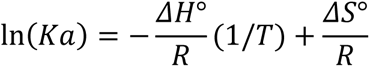

yields an Arrhenius plot [ln(Ka) as a function of 1/T] that displays a straight line, with a respective slope and y-axis intercept of 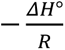 and 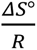, which afford Δ*H*° and Δ*S*°. Δ*G*° is calculated at 37 ℃.

### Molecular modeling

Molecular modeling was performed using the Molecular Operating Environment software (Chemical Computing Group, Montreal, Canada) and Amber94/99 force field parameters and a distance-dependent dielectric constant of ε = 4r (where r is the distance between two atoms) [55,56]. The initial structures of the d(AAGGG)_4_ and r(AAGGG)_4_ G4 conformations were constructed based on the crystal structure of the hybrid-type G4 (PDB ID: 7DJU) and d(TGGGT)_4_ G4 (PDB ID: 6p45) [57], respectively. Distance and dihedral-angle constraints were imposed on the guanine residues within the three G-tetrads during the process. The structure of the r(ACAGG)_11_ hairpin conformation was created using MC-Sym pipeline (University of Montreal, Montreal, Canada) [58] with default parameters and then further refined to yield the initial structure for use in the final minimization. The resulting structures were solvated by adding H_2_O molecules and Na^+^ ions, and the systems were minimized without constraints to the points where the root-mean-square gradients were <0.001 kcal/(mol Å).

## Data availability

All data needed to evaluate the conclusions in the paper are present in the paper. Additional data related to this paper may be requested from the corresponding author upon reasonable request.

## Acknowledgments

We thank editage for the English language review.

## Funding and additional information

This work was supported by the Japan Agency for Medical Research and Development (JP21ak0101131, JP23ek0109651, JP23ek0109591 to N.S.), Japan Society for the Promotion of Science KAKENHI (JP23H00373, JP22K19297 to N.S.), Japan Science and Technology Agency Fusion Oriented Research for disruptive Science and Technology Program (JPMJFR2043 to N.S.), program of the Joint Usage/Research Center for Developmental Medicine (IMEG, Kumamoto University), and program of the Inter-University Research Network for High Depth Omics (IMEG, Kumamoto University).

## Footnotes

### Abbreviations

CANVAS: cerebellar ataxia with neuropathy and vestibular areflexia syndrome
DMS: dimethyl sulfate
FAM: 6-carboxyfluorescein
FTD: frontotemporal dementia
FXTAS: fragile X-associated tremor/ataxia syndrome
IMEG: Institute of Molecular Embryology and Genetics
MOE: Molecular Operating Environment
RAN: repeat-associated non-AUG
RBP: RNA-binding protein
RFC1: replication factor C subunit 1
STR: short tandem repeat
TFO: triplex-forming oligonucleotide.

